# N2B27 media formulations influence gastruloid development

**DOI:** 10.1101/2025.03.15.643474

**Authors:** Tina Balayo, Sharna Lunn, Pau Pascual-Mas, Ulla-Maj Fiuza, Amruta Vasudevan, Joshua D. Frenster, Hannah Y. Galloon, Raquel Flores Peirats, Alfonso Martinez Arias, André Dias, David A. Turner

## Abstract

Gastruloids are 3D aggregates of pluripotent stem cells grown in suspension culture that mimic many aspects of gastrulation and early axial elongation. The N2B27 basal medium in which mouse gastruloids are cultured can either be home-made (HM-N2B27) with materials of known origin, or commercially sourced (NDiff227), where the exact formulation is unknown. In this study we examined whether these formulations resulted in significant differences in gastruloid development. Our results reveal that while both media enable the standard gastruloid elongation, HM-N2B27 results in gastruloids that start the elongation process earlier, have higher number of cells and an increased anterior domain. RNAseq analysis showed significant differences in cell fate specification, with HM-N2B27 gastruloids exhibiting higher expression of spinal cord-related genes, while NDiff227 favours mesodermal differentiation. Furthermore, differential gene enrichment analysis suggests that changes in key signalling pathways underline the differences between HM-N2B27 and NDiff227 gastruloids. These findings highlight the importance of basal media composition for gastruloid development, underscoring the need for careful media selection during *in vitro* engineering of stem cell-based embryo models.

**Summary statement:** In this work we explored the cellular and molecular sensitiveness of the mouse gastruloid model system to variations in N2B27 media formulations.

## Introduction

The development of an embryo is a highly organised progression of cell fate decisions which control the allocation of distinct cell types, and coordinate cell movements and tissue morphogenesis. Many of the mechanisms involved in early embryonic patterning have been derived from *in vivo* approaches, however in the case of mammals, especially human embryo development, there are a number of challenges, technical and ethical, which need to be overcome (Rossant and Tam, 2021; Rugg-Gunn et al., 2023).

*In vitro* model systems, making use of pluripotent stem cells (PSCs), provide an alternative and tractable experimental approach to study these developmental processes (Fu et al., 2021), and have become useful tools for understanding the process of cell fate decision-making in a physiological context. When grown in 3D and exposed to the appropriate signals, they can generate organoids which can be used to study the formation of specific tissue types and ‘organ-like’ structures (e.g. optic cup, gut, cerebral organoids) (Eiraku and Sasai, 2012; Sasai, 2013; Sasai et al., 2012). Gastruloids, embryonic organoids, are made from small numbers of aggregated PSCs (Turner et al., 2016; Turner and Martinez Arias, 2024). When they are exposed to a short pulse of Wnt/β-catenin signalling after 48h of culture in differentiation media, they mimic many aspects of the early gastrulating embryo such as symmetry-breaking with a polarisation of posterior-related gene expression, axial elongation, and the formation of all three embryonic axes with collinear expression of Hox-genes along the anteroposterior axis (Beccari et al., 2018; Dias et al., 2025; Turner et al., 2017; Turner et al., 2014a; van den Brink et al., 2014). A key feature of the gastruloid model system lies in its high reproducibility, which is critical to generate a deep understanding of developmental processes (Merle et al., 2024; Turner et al., 2017). It is therefore important to understand and properly characterise the factors that guide their differentiation, maintain their reproducibility, and importantly, acknowledge the variables that lead to suboptimal or alternative/variable phenotypes (Rosen et al., 2022). Recent work has shown that factors such as 2D culture conditions of the starting cell population (Blotenburg et al., 2024), initial number of cells (Bennabi et al., 2024; Fiuza et al., 2024; van den Brink et al., 2014), metabolism (Dingare et al., 2024; Luque et al., 2023; Stapornwongkul et al., 2023) and precision in the execution of the protocol (Dias et al., 2025) play a critical role in gastruloid variability and can have a significant impact on fate specification. A critical element of all these variables is the basal medium in which cells and gastruloids are grown: N2B27.

The N2B27 media was initially developed as a chemically-defined neural differentiation media (Mulas et al., 2019; Ying and Smith, 2003; Ying et al., 2003), however due to the minimal nature of its composition, it served as a useful base in which to add key signalling components to direct PSCs (either in monolayer or as gastruloids) to distinct lineages in a controlled manner (e.g. refs. (Gouti et al., 2014; Hennessy et al., 2023; Turner et al., 2014a; Turner et al., 2014b; Turner et al., 2014c; van den Brink et al., 2014)). N2B27 is primarily made from a 1:1 mixture of DMEM:F12 and Neurobasal media with added N2 and B27 (containing vitamin A) supplements (Mulas et al., 2019). A commonly used commercial version of this medium is known as NDiff®227 (NDiff227) and, presumably, contains similar components, although it is not possible to determine the exact quantities or makeup of this proprietary medium. Anecdotal evidence from several groups has suggested that gastruloid growth and development may differ depending on what medium is used: either homemade N2B27 or NDiff227 in the extreme. However, to our knowledge there has not been a systematic analysis of the differences between the two formulations in the context of gastruloid development.

Here, we undertook a short study across two laboratories to quantitatively assess the effect of using a homemade N2B27 media formulation (HM-N2B27) or NDiff227 on gastruloid development, morphology and gene expression. We find that although the shape of the gastruloids is broadly consistent, there are significant changes in morphometric variables, such as the length and elongation index of the gastruloid. Also, we detected differences in cell fates, with HM-N2B27 gastruloids showing an increased expression of spinal cord-related genes whereas NDiff227 gastruloids display higher levels of paraxial mesoderm-associated genes. A broader analysis on gene ontology (GO) and biological processes suggests that changes in key signalling pathways could underlie these observed differences in the gastruloids cultured with the two media. Overall, our work indicates that while distinct N2B27 media formulations do not change the reproducibility of the model, they can impact gastruloids at both the cellular and molecular levels and suggest that the choice of media should be a key consideration in the experimental design.

## Results and Discussion

### Distinct N2B27 media formulations impact gastruloid development, morphology, and cell number

To understand if different N2B27 differentiation media formulations affect the development of mouse gastruloids, we set up several independent comparative experiments, across two laboratories, using commercial NDiff227 and home-made N2B27 (HM-N2B27) and observed the morphology of wildtype E14Tg2A gastruloids at 120h following the standard 48-72h CHIR pulse. Although the classical elongation (Beccari et al., 2018; Turner et al., 2017; van den Brink et al., 2014) was obtained using both types of differentiation media, we noticed several clear differences regarding gastruloid morphology and the frequency of either single or multi axis gastruloids (**Fig. 1; Fig. S1**). In the first instance, we observed that in all replicate experiments, HM-N2B27 had approximately 60% single axis elongations (**Fig. 1A & B**), whereas NDiff227 had a much higher frequency of single axis elongations (>90%; *n* = 5 replicates; > 100 individual gastruloids; **Fig. 1A & B**). Also, we observed that HM-N2B27 were greater in overall size (area) and displayed a more pronounced elongation in comparison to NDiff227 gastruloids (**Fig. 1A,C**).

**Figure 1:**
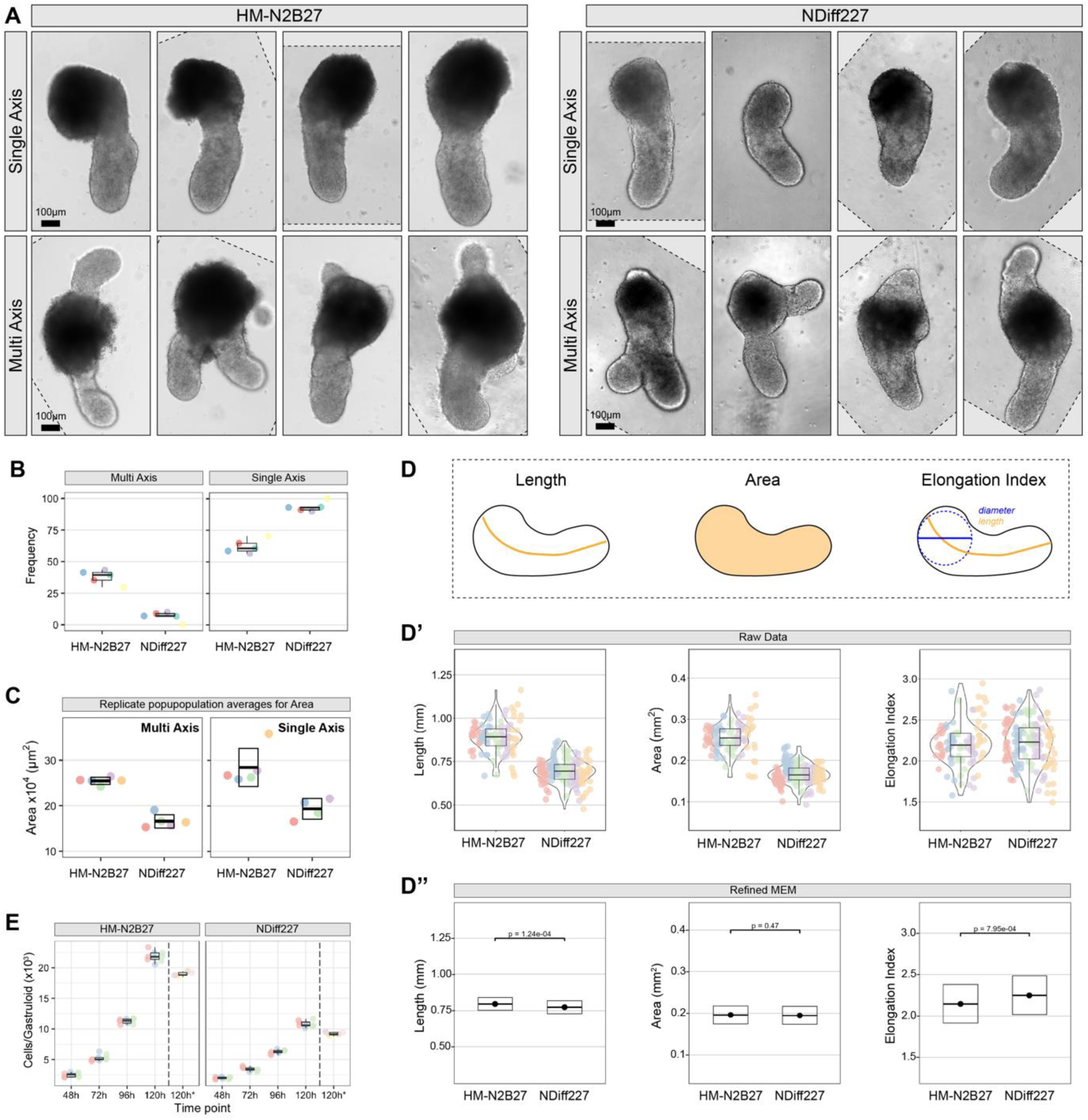
HM-N2B27 and NDiff227 media significantly affect gastruloid morphology and cell number. (**A**) Representative images selected from 5 independent experiments showing the general morphology of CHIR-treated gastruloids at 120h after aggregation, with either a single or multi axis grown in either E14Tg2A HM-N2B27 (left) or NDiff227 (right). (**B**) Replicate population averages for the frequency of single axis or multi-axes gastruloids in HM-N2B27 and NDiff227 conditions. (**C**) Replicate population averages for the area of HM-N2B27 or NDiff227 gastruloids (n = 5 for each replicate). (**D**) Morphometrics analysis of gastruloids in (A). Schematics indicate how each measurement was obtained. (**D’**) Raw quantification of the length, area, and elongation index of single axis gastruloids in HM-N2B27 or NDiff227. Individual gastruloids are represented as coloured circles, with distinct colours for each independent batch, and suggest an increase in the area and length of HM-N2B27 gastruloids in comparison to NDiff227 counterparts. (**D’’**) Predictive averages (±95% CI) generated from refined mixed effects models (MEM) of data from ≥95 gastruloids per media (see **Material and Methods**, **Fig. S2** and **Table S1**), highlighting significant differences in length and elongation index between HM-N2B27 and NDiff227. (**E**) Average number of cells in pooled E14Tg2A gastruloids from 3 independent experiments or from 5 independent gastruloids (120h*; right of hashed line) at the indicated timepoints. Data indicate a significant increase in the number of cells in HM-N2B27 gastruloids in comparison to NDiff227 counterparts. See supplemental Table S2 for significance testing for pooled gastruloids. A Mann-Whitney U test was used to compare the cell counts from individual gastruloids at 120h (120h*; W = 25; *p* = 0.0079). Scale bar indicates 100 μm.

A more thorough morphometrics analysis of the single axis gastruloids from both media was implemented, using mixed effects models (MEMs) that account for the combined influence of length, area and elongation index (**Fig. 1D & Fig. S2**; Lunn *et al.,* in preparation). HM-N2B27 gastruloids have a greater overall length (*p* = 1.24×10^−4^) whilst NDiff227 gastruloids have a greater elongation index (*p* = 7.95×10^−4^; **Fig. 1D, S2, & Table S1**). Although this may appear counterintuitive, these results can be explained by understanding the difference between the two measurements. Whereas length is the measure between the two furthest poles, the elongation index is calculated as a ratio between the diameter of the largest inscribed circle within the gastruloid and the gastruloid total length (**Fig. 1D**) (Guiet et al., 2021). This distinction is exemplified upon observation of the gastruloids: those developed in HM-N2B27 have a larger anterior pole relative to their posterior compartment, which contributes to an increased overall length but reduces its elongation index. On the other hand, NDiff227 gastruloids display a more consistent shape from pole to pole, which results in a larger elongation index (**Fig. 1A,D, Fig. S2, & Table S1**). Although the raw area data of single axis gastruloids is significantly different between the two formulations, once the influence of other variables was mitigated, this difference was attributed to the length, elongation index and their interactions with each other (**Fig. 1, Table S1**). Measurements of the overall area of multi-axis gastruloids also indicate that the use of HM-N2B27 media can result in gastruloids with a greater area than NDiff227 (*p* < 0.005; **Fig. 1C**). However, in this case the length and elongation index of multi axis gastruloids could not be determined as the longest axis to measure was often ambiguous.

The observed size difference between gastruloids grown in HM-N2B27 or NDiff227 led us to consider whether different media conditions directly influenced gastruloid cell number. To examine this, we quantified the number of cells in gastruloids from both culture media over time. Due to the limited number of cells at early time points, multiple gastruloids were pooled together to improve count accuracy (see **Material and Methods** section for details). At each time-point, with the exception of 48h, HM-N2B27 gastruloids showed consistent and significantly greater numbers of cells than NDiff227 gastruloids (**Fig. 1E, Table S2**; *p* < 0.0001). As averaging whole pools of gastruloids has the potential to introduce averaging errors, the cells from individual gastruloids (*n* = 5) were also counted at the 120h time-point (**Fig. 1E**; denoted 120h*) and the results indicate that HM-N2B27 gastruloids have a significantly higher, roughly double, number of cells than NDiff227 (**Fig. 1E**; Mann-Whitney test, *W* = 25; *p* = 0.0079). As several reports showed that increasing the number of seeding cells results in multi axis formation (Bennabi et al., 2024; Fiuza et al., 2024; van den Brink et al., 2014), it is possible that the higher frequency of multi axis in HM-N2B27 gastruloids is linked to the increased number of cells in comparison to NDiff227 gastruloids. However, as the number of seeding cells was kept consistent in these experiments, our results suggest that the use of HM-N2B27 or NDiff227 media significantly impacts the rate of cell proliferation, survival and/or differentiation during gastruloid development. In agreement, a recent study suggested that cell competition was reduced in gastruloids developed with HM-N2B27 media in comparison to NDiff227 (Frenster et al., 2024).

A further morphological difference was also found which related to the timing of gastruloid development. NDiff227 gastruloids usually displayed a round shape at 96h, though some ellipsoid and ogival morphologies were also found depending on the batch. In contrast, HM-N2B27 gastruloids displayed more pronounced ellipsoid and ogival shapes at this time (**Fig. 2** and **S3**). This suggests that the process of elongation started earlier in the homemade condition. In agreement, we noticed some batch-to-batch variability in the commercial media conditions, NDiff227 gastruloids usually displayed a round shape at 96h whereas the majority of HM-N2B27 gastruloids had already started the elongation process at this time (**Fig. 2** and **S3**). Concomitantly, we observed that the cadherin switch from Cdh1 (E-cadherin) to Cdh2 (N-cadherin) (Basilicata et al., 2016; Mayran et al., 2023), started around 6h earlier with 90h HM-N2B27 gastruloids showing an expression pattern more similar to 96h NDiff227 gastruloids (**Fig. 2**). Together, these results suggest that distinct N2B27 media formulations can impact on gastruloid developmental time, size and morphology, particularly affecting the development of the anterior compartment and the length of the posterior, classical elongation.

**Figure 2:**
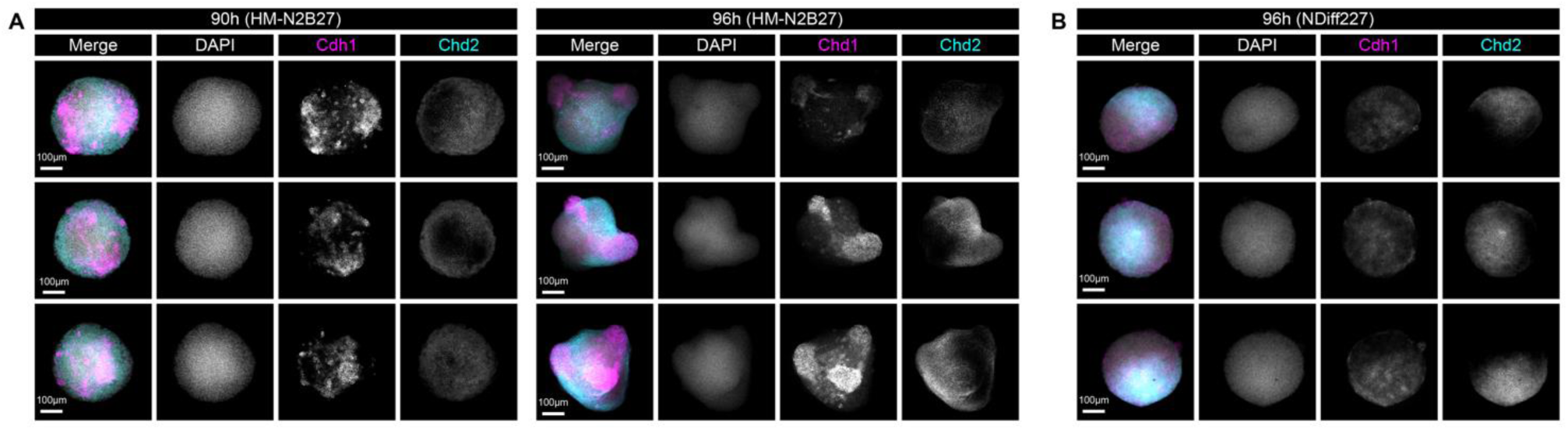
NDiff227 gastruloids display a developmental delay in comparison to HM-N2B27 gastruloids. Immunofluorescence staining for Cdh1 (E-cadherin), Cdh2 (N-cadherin) and DAPI, showing that 96h E14Tg2A NDiff227 gastruloids (**B**) display a more similar cadherin pattern and overall morphology (see also **Figure S3**) in comparison to 90h E14Tg2A HM-N2B27 gastruloids, than to their 96h counterparts (**A**). In composite images Cdh1 is shown in magenta, Cdh2 in cyan, and DAPI in grey. Scale bar indicates 100 μm.

### N2B27 media influences key signalling networks and cell fate specification in the gastruloid model system

To understand whether the differences in gastruloid development we observed in different media formulations were also reflected at the level of cell fate specification, we harvested gastruloids at 120h and analysed their gene expression by bulk RNA sequencing (RNAseq; **Fig. 3**). Statistical pairwise comparison showed that 196 genes were differentially expressed (Log fold change (LFC) = 1.5, *p*-value threshold ≥ 0.05) between HM-N2B27 and NDiff227 samples (**Table S3, Fig. 3A**). Principal component analysis (PCA) revealed that both samples form distinct, separate clusters in the first dimension (accounting for 71% of the variance), with a significant distance between the two (**Fig. 3B**). The top 100 genes based on loadings of PCA dimension 1 and 2 (**Table S4**, **Fig. 3B**) suggest that these differences might be associated with fate specification, particularly with different amounts of neuroectoderm (e.g. *Pax6* and *Pou3f2*) and presomitic mesoderm (e.g. *Tbx6*, *Lfng* and *Mesp2*). Although both gastruloids made either with HM-N2B27 or NDiff227 developed the classical derivatives of the late/posterior PS (Dias et al., 2025), an in-depth analysis of specific fate markers indicated significant differences in the levels of genes associated with neuroectoderm and paraxial mesoderm. HM-N2B27 gastruloids exhibited higher levels of *Sox1*, *Sox2*, *Sox3* and *Pax6* than those made with NDiff227 (**Fig. 3C**). These differences in neuroectoderm are associated with spinal cord-like cells and not with brain-like structures (Pfister et al., 2007; van den Brink et al., 2020) as there was no significant expression of *Hesx1*, *Pou3f1* and *En1* (**Fig. 3C**). On the other hand, gastruloids made with NDiff227 expressed higher levels of *Tbx6*, *Dll1*, *Pcdh19*, *Lfng*, *Tcf15* and *Raldh2* (*Aldh1a2*) than those made with HM-N2B27 (**Fig. 3C**). These differences are associated with an increase in early/nascent, presomitic and paraxial mesoderm-like tissues (Dias and Aires, 2020; van den Brink et al., 2020). Interestingly, we found that several Notch-related genes, such as *Notch1*, *Dll1,* (Geffers et al., 2007), *Dll3* (Chapman et al., 2011), *Hes7* (Kageyama et al., 2007), *Mesp2* (Barrantes et al., 1999; Takahashi et al., 2000), *Runx1* (Li et al., 2018; Zhou et al., 2022) and *Nrarp* (Krebs et al., 2012; Lamar et al., 2001) were differentially regulated between HM-N2B27 and NDiff227 gastruloids at 120h (**Fig. S4**). Given the central role that Notch plays in the differentiation of mammalian axial progenitors towards the neural and mesodermal lineages (Cooper et al., 2024; French et al., 2024; Gray and Dale, 2010), it is possible that Notch signalling activity might vary depending on the N2B27 media used and that it drives the observed differences in neural and mesodermal-related gene expression between the two conditions. Testing this hypothesis would require further mechanistic studies involving systematic gain and loss of function experiments, along with the monitoring of both Notch signalling activity and the expression of neural and mesodermal marker genes. Finally, we also found no differences in lateral/intermediate mesoderm (e.g. *Lhx1*, *Osr1* and *Pax2* (Beccari et al., 2018; Dias et al., 2020; Dressler, 2009), endoderm (e.g. *Sox17*, *Foxa2 and Apela* (Beccari et al., 2018; Nowotschin et al., 2019; Pfister et al., 2007) or notochord-related gene expression (Beccari et al., 2018; Wymeersch et al., 2019) (e.g. *Noto and Shh*; **Fig. 3C**).

**Figure 3:**
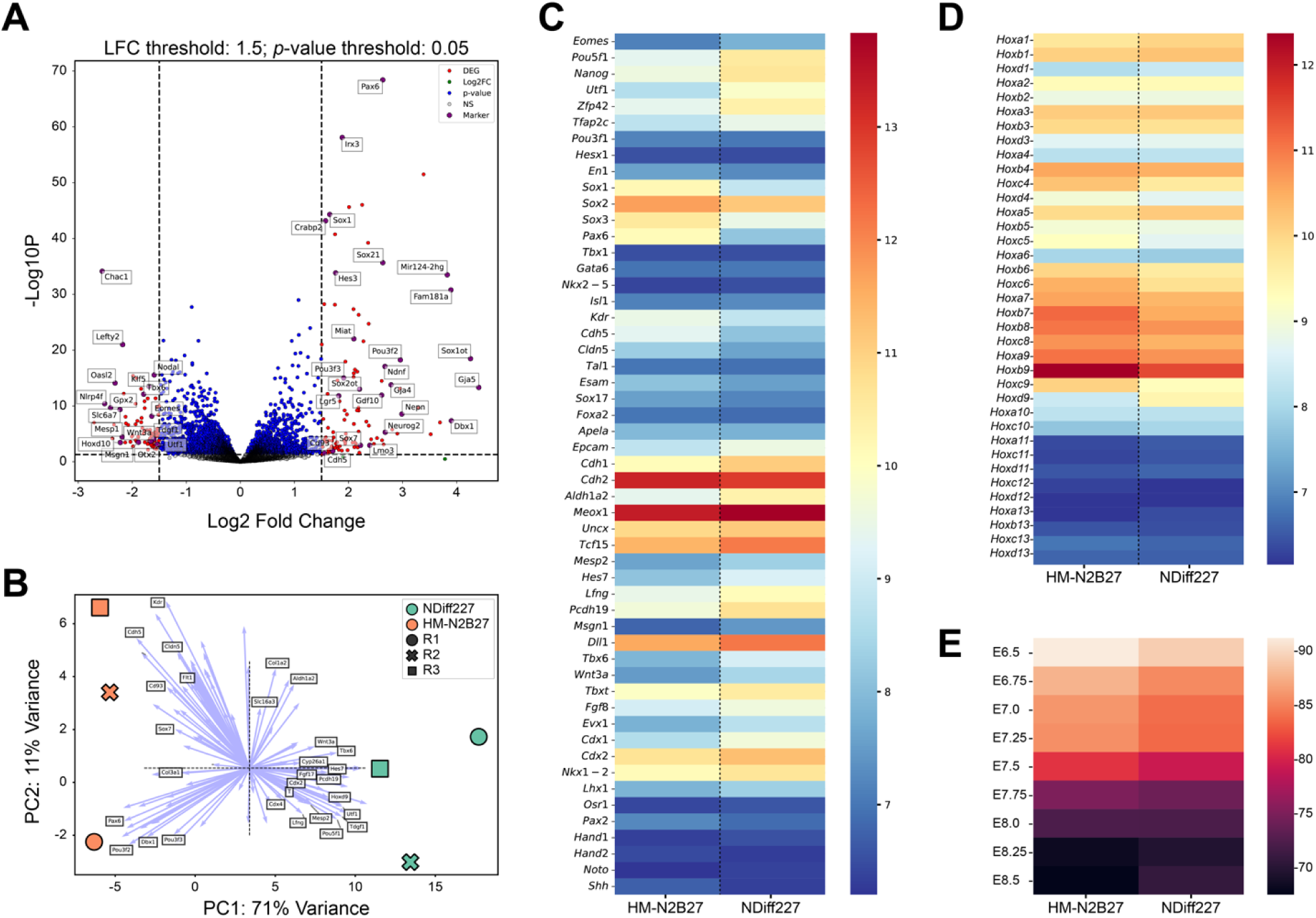
Transcriptomic analysis between HM-N2B27 and NDiff227 gastruloids. Bulk RNAseq analysis of E14Tg2A gastruloids harvested at 120h, cultured in either HM-N2B27 or NDiff227. (**A**) Volcano plot of differentially expressed genes (DEG) with a log-fold change (LFC) threshold of 1.5 (denoted by red), and significance threshold ≥ 0.05 (blue and red). Specific DEGs of interest are highlighted, and the full list can be found in Table S3. (**B**) Principal component analysis (PCA), with some of the top 100 genes based on loadings of PCA dimension 1 and 2 loadings (blue arrows; full gene list can be found in Table S4), highlighting the differences between NDiff227 (in green) and HM-N2B27 gastruloids (in orange; 3 replicates per condition). (**C**) Heatmap of selected fate marker genes (mean of the replicates variance-stabilized read counts), highlighting the differences and similarities between HM-N2B27 and NDiff227 gastruloids at 120h. Also, there is a difference in the expression of spinal cord (*Sox1*, *Sox2*, *Sox3* and *Pax6*) and paraxial mesoderm-related genes (*Aldh1a2*, *Meox1*, *Uncx* and *Tcf15*), between the two media conditions. Genes related to axial progenitors (e.g. *Tbxt*, *Cdx2* and *Nkx1-2*) are upregulated in NDiff227 gastruloids (**D**) Comparative heatmap (mean of the replicates variance-stabilized read counts) showing minor differences related to *Hox* gene expression. (**E**) Euclidean distance analysis between gastruloids cultured in the two media conditions and mouse embryos, from embryonic day (E) 6.5 to E8.5, highlights a similar temporal development between the two.

Euclidean distance analysis between the two types of gastruloids and mouse embryos (Pijuan-Sala et al., 2019) indicated that the development of the gastruloids occurred as expected, reaching early stages of axial elongation (Beccari et al., 2018), with gastruloids made with NDiff227 being more related to E8.25 mouse embryos whereas those obtained with HM-N2B27 seem similar to E8.5 embryos (**Fig. 3E**). Concomitantly, the expression of *Hox* genes was found to not differ substantially between the two types of gastruloids (**Fig. 3D**). Small differences were observed in *Hox* genes that at these stages are known to extend up to more anterior regions of the body axis (*Hoxc4*, Hoxc5, *Hoxc6* and *Hoxb9*) (Deschamps and Duboule, 2017; Goh et al., 1997; Minguillon et al., 2012; Nishimoto et al., 2014). These *Hox* genes were slightly upregulated in HM-N2B27 gastruloids (**Fig. 3D**), suggesting they might contain more anterior-like structures. In agreement, we found that some genes associated to endothelial cells were upregulated in gastruloids made with HM-N2B27 (e.g. *Kdr* and *Cdh5*, **Fig. 3C**). On the other side, NDiff227 gastruloids displayed higher levels of *Hoxd9*, which is more restricted to the caudal part of embryo, and genes like *Tbxt, Nkx1-2, Cdx2* and *Cdx1* (**Fig. 3C&D**), that are associated with the neuromesodermal competent population (Binagui-Casas et al., 2021). The increase of *Hoxa5* in these gastruloids (**Fig. 3D**) is likely due to the increase in mesodermal-like structures (Minguillon et al., 2012). Another key difference between NDiff227 and HM-N2B27 gastruloids is that the former contains significantly higher levels of *E-cadherin* (*Cdh1)*, *Oct4* (*Pou5f1*), *Nanog*, *Utf1*, *Rex1* (*Zfp42)* and *Tfap2c* (**Fig. 3C**). These are genes that have been related to both primordial germ-like cells (PGCs) (Cooke et al., 2023; van den Brink et al., 2020) and ectopic pluripotency (Suppinger et al., 2023) at this stage in mouse gastruloids.

A broader analysis focused on signalling pathways, gene ontology, and biological processes also revealed significant differences between HM-N2B27 and NDiff227 gastruloids at 120h. In addition to an enrichment of pluripotency and mesoderm-related processes, our results suggest that the expression of genes related to the FGF, p53 and TGFβ/Nodal signalling pathways is enhanced in gastruloids made with NDiff227 (**Fig. 4A; Table S5**). For instance, the enrichment of the latter is a consequence of the upregulation of both Nodal and its downstream targets, *Lefty1* and *Lefty2* (Bisgrove et al., 1999; Meno et al., 1999) in NDiff227 gastruloids (**Fig. S4**, **Table S5**). In contrast, PI3K-Akt, cGMP-PKG, Ras, and Hedgehog signalling were enriched in HM-N2B27 gastruloids (**Fig. 4A**, top; **Table S5**). The PI3K/Akt signal cascade is a known regulator of several key cellular functions such as growth, migration, differentiation and cell survival (Riley et al., 2005). Recent work in gastruloids has shown that this signalling pathway is required for normal axis elongation (its inhibition reduces gastruloid length) and can induce proliferative activity in the anterior compartment of gastruloids, leading to its expansion (Underhill and Toettcher, 2023). These results are consistent with our observations as HM-N2B27 gastruloids have an enrichment of PI3K/Akt-related genes (e.g. *Igf1*, *Ccnd2*, *Thbs1* and *Itgb8*) (Scrima et al., 2012; Shu et al., 2019; Stitt et al., 2004; Tuoheti et al., 2024) and display an increased anterior compartment and length in comparison to NDiff227 gastruloids (**Fig. 1** and **S4, Table S5**). Finally, our GO analysis also suggested an enrichment of cellular processes related to cell adhesion/motility, extra-cellular matrix and cytoskeleton regulation with HM-N2B27 (**Fig. 4A, Table S5**) due to the upregulation of *Mylk* (*Mlck*) (Blaser et al., 2006; Cai et al., 1998; Samson et al., 2019; Weiser et al., 2009), *Nrcam* (Grumet et al., 1991), *Lama4* (Malinda and Kleinman, 1996)*, Col15a1* (Eklund et al., 2001; Rasi et al., 2010), *Col3a1* (Kuivaniemi and Tromp, 2019), and *Thbs1* (Murphy-Ullrich, 2019) in HM-N2B27 gastruloids (**Fig. S4, Table S5**). In various studies (Cermola et al., 2022; de Jong et al., 2024; Fiuza et al., 2024; Mayran et al., 2023; Pineau et al., 2025; Veenvliet et al., 2020) these processes have been shown to be critical for gastruloid elongation. For instance, forced reduction in actin polymerization through the use of cytochalasin D between 90-120h resulted in a more prominent elongation in mouse gastruloids treated with CHIR (Fiuza et al., 2024). Therefore, our work provides an extensive list of pathways and target genes that can be used in subsequent studies aimed at identifying and modulating the factors that play a role during axial elongation, both *in vivo* and *in vitro*. As the gastruloid model system is not confined to mouse ESCs, and human gastruloids have recently been developed (Moris et al., 2020), it will be interesting in the future to address whether the human system is also sensitive to distinct media formulations and to understand what the biological implications of such results.

**Figure 4:**
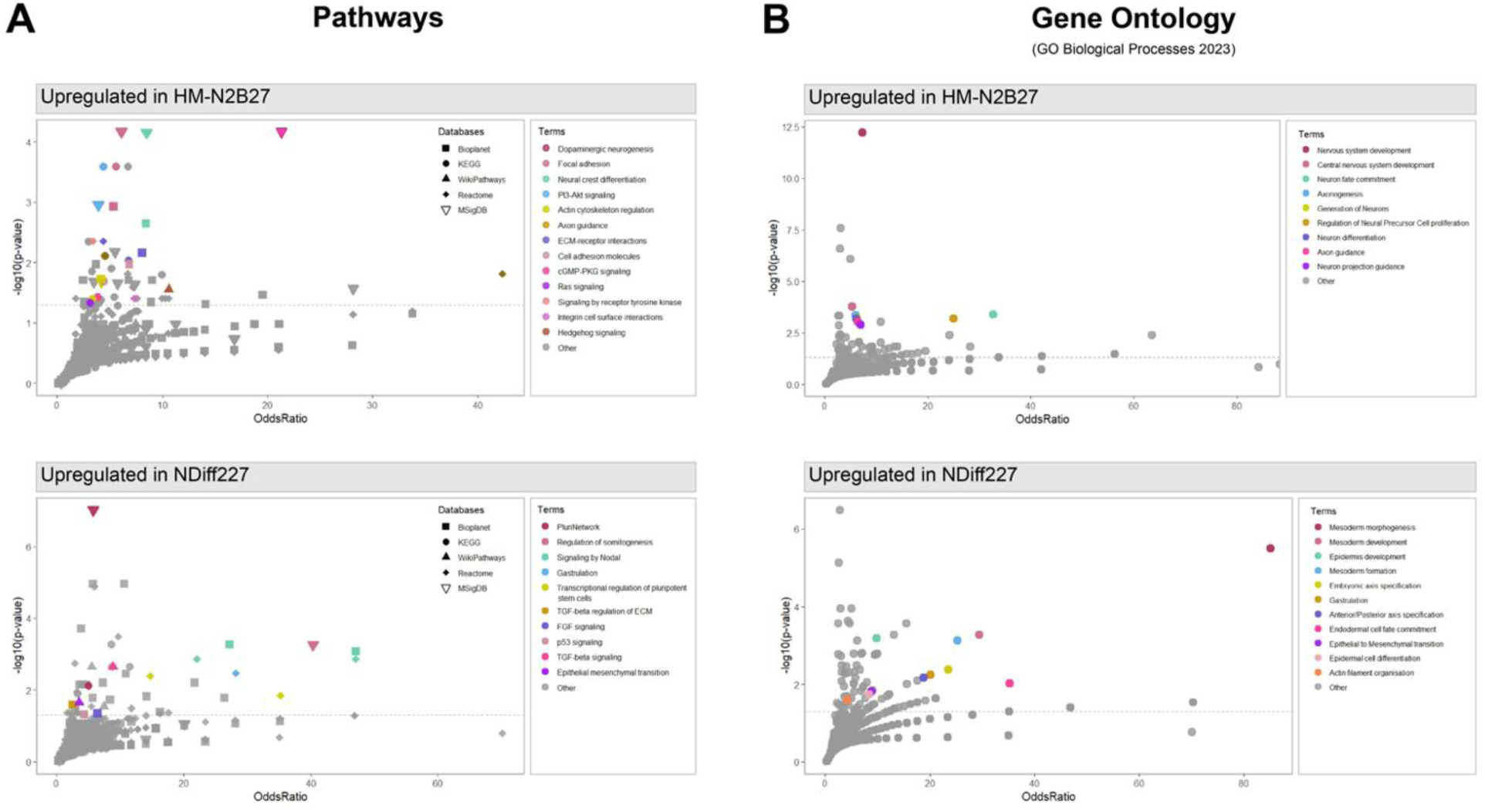
N2B27 and NDiff227 media differently impact key signalling pathways and biological processes during gastruloid development. Over-representation analysis (ORA) conducted on the DEGs obtained from the bulk RNAseq of HM-N2B27 and NDiff227 E14Tg2A gastruloids at 120h, highlighting differences in signalling pathways (**A**) and biological processes (**B**) between the two media conditions. All datapoints represent individual annotation terms from Pathways databases (**A**), with each shape pertaining to a unique database (refer to legend ‘Databases’), and the GO Biological Processes 2023 database (**B**). The significantly enriched and biologically meaningful datapoints are coloured according to the annotation terms. The x-axis is the Odds Ratio, the y-axis is −log10(p-value adjusted for multiple testing), and the dotted line corresponds to an adjusted p-value of 0.05.

## Conclusion

Our findings, independently developed in two laboratories, demonstrate that variations in N2B27 media formulations can significantly influence key aspects of mouse gastruloid development. These include gastruloid morphometrics, developmental timing, cell fate specification, signalling networks and the rate of cell proliferation, survival and/or differentiation. Therefore, we propose that the N2B27 media (and other similar culturing media) should be considered an important experimental variable in gastruloid studies and, more broadly, in the *in vitro* engineering of stem cell-based embryo models.

## Limitations

Due to the specific nature and scope of this study, there are limitations that a more in-depth, mechanistic, analysis could address in the future. One limitation is the lack of assessment of inter- and intra-batch transcriptomic variability. Addressing this question would require single-cell RNA sequencing, which is costly and unlikely to provide substantial additional insights to our work. Another limitation has to do with understanding whether, and if so how, the cellular/morphological changes in gastruloids cultured in the two media are linked to the observed genetic differences. This would require further systematic work beyond the question we set out to evaluate in this study - determining whether commercial and homemade N2B27 media formulations function similarly in mouse gastruloid development. A third limitation regards the lack of assessment on whether the activity of the potential differentially regulated signalling pathways was indeed affected by the two media compositions and whether the differentially expressed genes associated with processes such as cell adhesion/motility, extracellular matrix organization, and cytoskeletal regulation are functionally relevant for mouse gastruloid development. Addressing these questions would require specific reporter cell lines, which are unavailable in our laboratories, as well as mutagenesis or pharmacological treatments during gastruloid development. While we aim to conduct some of these experiments in the future, they fall beyond the scope of this work. Finally, another limitation of our study relates to our inability to properly compare the ingredients of the two N2B27 media compositions due to the proprietary nature of the commercial NDiff227 recipe.

## Materials and Methods

### N2B27 media formulations

Commercial NDiff®227 (NDiff227) was obtained from Takara Bio (Takara, Y40002) and the following lots were used in this study: ‘AM30020S’, ‘AM10016S’, ‘AM90020S’, ‘AN30020S’ and ‘AO30013S’. NDiff227 media displaying a non-uniform colour is not optimal for gastruloid culture and, therefore, was not used in this study. Home-made N2B27 was prepared in lots of 50ml by mixing equal volumes of freshly made N2 and B27 media, together with β-mercaptoethanol (Invitrogen; 31350010) at a concentration of 1:1000. 25ml of N2 media were prepared using 24.5ml of DMEM/F12 (1:1) (Gibco, 21331-020), 250µl of N-2 Supplement (100x; Gibco, 17502-048; Lot: 2584689, 2584685 and 2831191) and 250µl of L-Glutamine (Invitrogen, 25030-024). 25ml of B27 media were prepared using 24.25ml of Neurobasal (Gibco, 21103-049), 500µl of B-27 Supplement (50x) (Gibco, 17504-044; Lot: 2596518, 2596510 and 2814925) and 250µl of L-Glutamine. Home-made N2B27 used to develop gastruloids from mouse ESCs pre-cultured with 2iLIF media (2iL, see below) was prepared in the Liverpool laboratory (standard was prepared in Barcelona), following a similar recipe but some ingredients were acquired from different suppliers, batches, or catalogue numbers: DMEM/F12 (1:1) (Gibco; 11320074), N-2 supplement (Gibco; 175020-48; Lot: 2868549, and 2868551), B-27 supplement (50x) (Gibco, 17504-044; Lot: 2814935, 2814937, and 2831197), and Glutamax (ThermoFisher, 35050038) was used instead of L-Glutamine. The N-2 Supplement, B-27 supplement, and L-Glutamine were kept at −70°C and taken to 4°C only the day before the preparation of HM-N2B27. Similarly to NDiff227, HM-N2B27 was also protected from light and stored at 4°C for no more than 2 weeks. Both types of N2B27 media were batch-tested before the experiments described in this study to assess their ability to generate gastruloids and the degree of morphological inter- and intra-batch variation.

### Cell lines and gastruloid development

E14-Tg2A (Hooper et al., 1987) mouse embryonic stem cells (ESCs) were maintained in 10% (standard) or 15% (high serum condition) ESLIF (ESL) media and used for gastruloid development, 300 cells per gastruloid, as indicated in Dias et al 2025 (Dias et al., 2025). Standard E14-Tg2A gastruloids were developed in non-adherent U-bottom 96-well plates (Greiner, 650185) whereas cells pre-cultured with higher levels of serum were aggregated for gastruloid formation in Ultra-low attachment plates (Greiner, 650970). The Bra::GFP (Fehling et al., 2003) and E14-Tg2A ESC lines pre-cultured for at least 2 weeks in 2iL (N2B27 base medium supplemented with 3 μM CHIR9901 (Tocris; 4423), 1 μM PD03 (Tocris,) and 11ng/mL of LIF (Qkine, Qk018) were used to make gastruloids as previously described (Beccari et al., 2018; Turner et al., 2017; van den Brink et al., 2014) with the following modifications. One passage before plating, cells were sub-cultured into a 25 cm^2^ tissue-culture flasks in 5ml 2iL with a medium change the following day. On the day of gastruloid plating (0 hours (h), after aggregation; AA), the outer wells of a sterile, non-tissue-culture-treated U-bottomed 96-well plate were filled with 100 μL of PBS to minimise evaporation of the gastruloid culture medium. Cells were washed with 3 ml PBS^−/−^ and incubated with 2 mL TrypLE for no longer than 3 minutes in a 37°C 5% CO_2_ incubator. TrypLE was diluted with 7 mL growth media, and cell colonies were dissociated by repeated up-and-down pipetting up for ∼5 times across the growth surface. The cell suspension was transferred to a 15 mL centrifuge tube and cell pelleted at 200g for 5 minutes. The supernatant was aspirated, cells washed in warm 10 mL PBS^+/+^, and centrifuged at 200g for 5 minutes, repeated twice, to ensure minimal carryover of PBS/TrypLE. The cell pellet was covered in 2 mL N2B27 and fully resuspended using a p1000 pipette to ensure a single-cell suspension is obtained with minimal bubbling. Cells were counted using a BioRad TC20 automated cell counter and 10 µL of cell suspension. The required number of cells (between 200-300 cells/well) was determined and scaled up to generate a 5 mL suspension per plate. To each well not containing the PBS evaporation buffer, 40 µL of the cell suspension was added using a multichannel pipette and left to aggregate for 48 hours in a 37°C; 5% CO_2_ incubator. At 48h AA, gastruloids were stimulated with 150 µL N2B27 supplemented with 3 μM CHIR. The next day (72h AA) gastruloids were “washed” and then fed with fresh N2B27 (i.e. 150 µL of N2B27 was removed and replaced with 150 µL of fresh N2B27 twice). At 96h AA, 150 µL N2B27 was replaced with fresh media. Every addition of N2B27 between and including 48h AA and 96h AA was administered with vigorous agitation to ensure limited attachment of gastruloids to the well surface. Images of gastruloids were taken using an Olympus brightfield CKX53 microscope with an EP50 camera on Zeiss Cell Observer fluorescence microscope (5x or 10x lenses depending on the size of the gastruloid) or a Nikon Ti-E inverted widefield microscope (set to 37°C, 5% CO_2_) using a 10x 0.3 NA objective with data capture using the Nikon NIS-Elements AR software. Gastruloids that did not elongate were removed from the analysis: around 10% with HM-N2B27 and 20% with NDiff227, depending on the lot used.

### Gastruloid cell counting

E14-Tg2A gastruloids grown in HM-N2B27 or NDiff227 were pooled, washed, and used for cell counting (Countess 3 Cell Counters, Invitrogen) at different stages according to the following quantities: 48h: 96 gastruloids; 72h:48 gastruloids; 96h:24 gastruloids; 120h:12 gastruloids. Dissociation was done with Accutase (Capricorn Scientific, ACC-1B), in accordance with Dias et al 2025 (Dias et al., 2025). Three independent experiments were performed and 5 gastruloids for each condition at 120h were also dissociated and their cells counted in a separate manner. To avoid any bias/difference between the two N2B27 media conditions during aggregation, temporal single gastruloid counts were also performed, with similar results, in gastruloids developed in ultra-low attachment plates (data not shown).

### Statistical analysis

Statistical analysis and graphs were produced using Prism Software (GraphPad) & RStudio (Team, 2019). Unpaired, parametric t-tests and two-way ANOVAs with Tukey post-hoc comparisons were used to analyse cell count (**Fig. 1**). Mixed-effects models used for morphological analysis (**Fig. 1**) considered media, area, elongation index and length as fixed variables, whilst batch replicates were included as a random effect in a random intercept model, and random slope models were used where appropriate. Each model was optimised to fit statistical assumptions including homoscedasticity and normal distribution of residuals; performance of each model can be found in Supplemental **Fig. S2**. After accommodating for interactions between fixed variables, Tukey post-hoc pairwise comparisons were used to determine statistical significance. Multi-axis gastruloid area was compared using an unpaired Mann-Whitney test (**Fig. S2**).

### Immunofluorescence staining and imaging

Immunofluorescence stains were done in accordance with Fiuza et al., 2024 (Fiuza et al., 2024). The primary antibodies used were goat anti-E-Cadherin (AF648; R&D Systems; 1:500) and rabbit anti-N-Cadherin (ab18203; Abcam; 1:200). The secondary antibodies used were donkey anti-rabbit Alexa Fluor 647 (A31573; ThermoFisher; 1:500) and donkey anti-goat Alexa Fluor 488 (A11055; ThermoFisher; 1:500). Imaging was performed using a Zeiss LSM980 confocal microscope with a 20x/0.8NA Plan-Apochromat objective. Data analysis was performed in the ImageJ package Fiji (Rueden et al., 2017; Schindelin et al., 2012).

### Bulk RNA sequencing and GO term analysis

Three independent experiments were done to generate three 96-well plates of E14-Tg2A gastruloids made with either HM-N2B27 or NDiff227 (48 gastruloids per sample, per plate). RNA from gastruloids was extracted using the Qiagen RNeasy Micro Kit (74004) and its concentration and purity were determined using PicoGreen and Fragment Analyzer (Agilent). Library preparation was done by the CRG Genomics Facility (Spain), using the TruSeq stranded mRNA Library Prep (ref. 20020595, Illumina) according to the manufacturer’s protocol. Sequencing was performed, also at the CRG Genomics Facility, in NextSeq 2000 and generated around 30M paired-end reads per sample. The preprocessing and downstream analysis of the bulk RNAseq data was done in accordance to Dias et al 2025 (Dias et al., 2025). Genes with an adjusted p-value < 0.05, determined via the Benjamini-Hochberg method for multiple testing correction, and an absolute log2 fold change > 1 were classified as significantly differentially expressed (DEGs). Over-representation analysis (ORA) of the DEGs was conducted using enrichR (version 3.2) and validated using WebGestalt 2024 (**Table S5**). The custom background gene list used for ORA was chosen to include all genes with non-zero counts from the bulk RNAseq analysis (**Table S5**).

## Acknowledgements

We would like to thank Jenny Nichols and Carla Mulas for advice and discussions regarding N2B27, Anna Bigas for the E14-Tg2A cell line, and Gordon Keller for the Bra::GFP cell line (Fehling et al., 2003). We are indebted to the University of Liverpool’s (UoL) Centre for Cell Imaging (CCI) facility for provision of state-of-the-art imaging equipment funded by the BBSRC (BB/R01390X/1) to the University of Liverpool’s CCI, and to the CRG Genomics and Imaging Units DAT was funded in this work by the BBSRC through a *New Investigator Grant* (BB/X000907/1; SL & AV), a *Strategic Longer and Larger* (sLoLa) Programme Grant (BB/Y00311X/1; HYG), and an NC3Rs PhD studentship (NC/T002131/1; HYG). AMA lab was funded by an ERC Advanced Grant (MiniEmbryoBlueprint 834580) and by the “Maria de Maeztu” Program for Units of Excellence in R&D (Grant No. CEX2018-000792-M). JDF was funded by a “la Caixa” Foundation (ID 100010434) Junior leader fellowship project (LCF/BQ/PI23/11970017). AD (ALTF 948-2022) and JDF (ALTF 605-2022) were funded by EMBO Postdoctoral Fellowships.

## Author Contributions

Conceived the study: TB, AMA, AD and DAT. Preliminary investigations: TB, RFP, JDF, AD, HYG, DAT, SL. Standard E14-Tg2A gastruloid experiments: TB, AD and RFP. Gastruloid experiments from 2iL-cultured Bra::GFP and E14-Tg2A ESCs: DAT, SL and JDF. Immunofluorescence staining: UMF. Bioinformatics and mathematical analyses: PPM, SL, AV, AD, and DAT. Original draft: AD and DAT. All authors edited the final manuscript.

## Data and resource availability

Raw and processed RNA sequencing data was submitted to Biostudies/ArrayExpress with the following accession information: “E-MTAB-14893”. The bioinformatics analysis pipeline, including all code and parameters, is available on the following GitHubs: https://github.com/stembryo-lab/HM-N2B27_vs_NDiff227_gastruloids and https://github.com/gastruloids/N2B27_vs_NDiff.

## Supplemental Figures

**Supplementary Figure 1:**
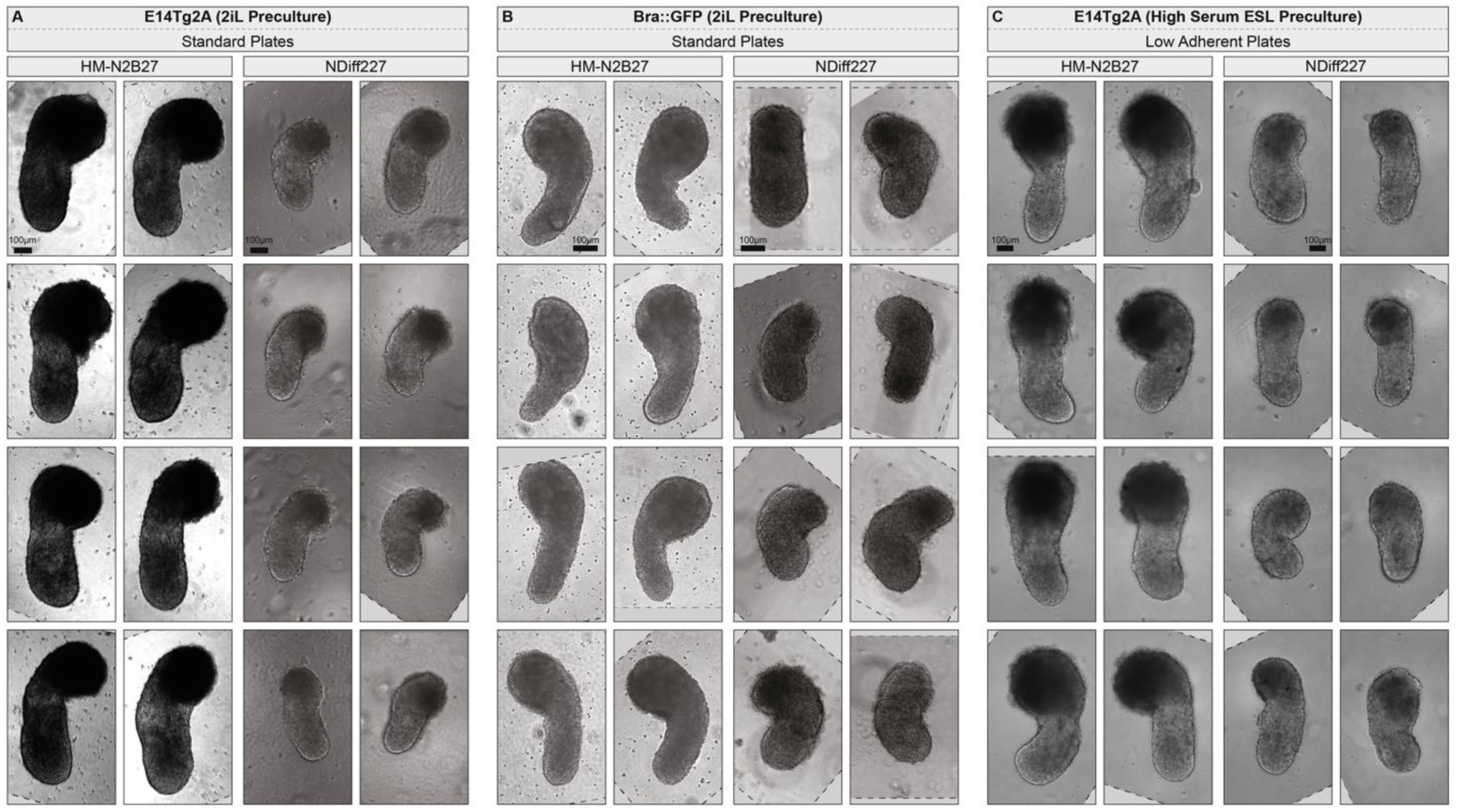
HM-N2B27 and NDiff227 gastruloids show consistent phenotypic differences across cell types, culture conditions, and laboratories. Example images of 120h single-axis gastruloids generated in different laboratories (see **Material and Methods**) with either E14Tg2A (A and C) or Bra::GFP ESCs (B) using different batches of both NDiff227, and HM-N2B27 raw materials, and different pre-culture conditions. Gastruloids in panels A and B were grown in standard 96-well plates and generated from cells pre-cultured for at least 2 weeks in 2iL pluripotency media. The gastruloids in panel C were formed in low-adherent 96-well plates and generated from cells cultured in ESL media with a high concentration of serum (see **Material and Methods**). HM-N2B27 consistently produce larger and longer gastruloids than NDiff227, regardless of the pre-culture conditions, or the 96-well plates used. Scale bar represents 100 μm.

**Supplementary Figure 2:**
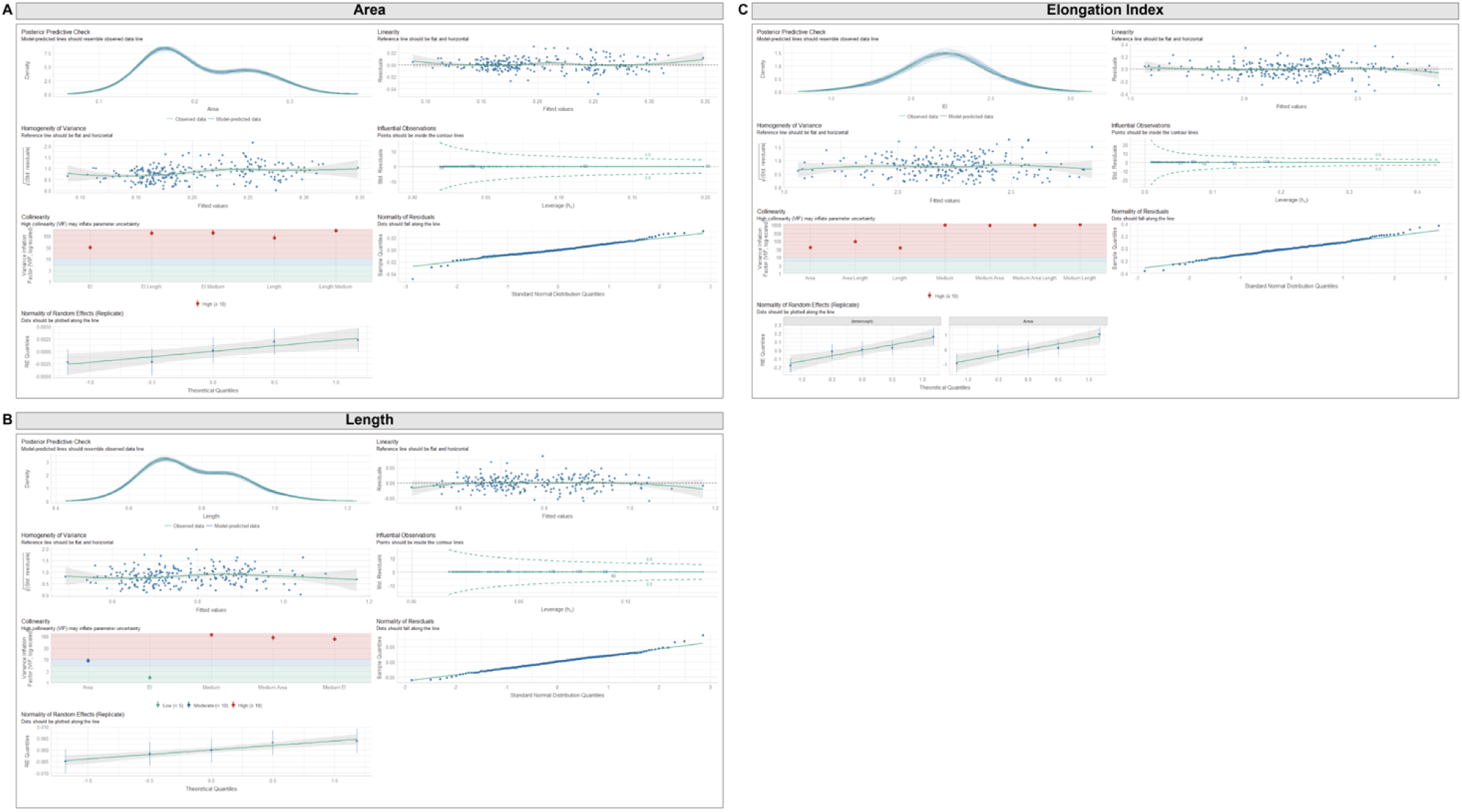
Performance analysis of mixed effects model to validate statistical conclusions made for elongation index (A), area (B) and length (C) of gastruloids in Figure 1D. Mixed-effects model used was as follows “lmer(EI ∼ MediumAreaLength + (Area|Replicate)) “, “lmer(Area ∼ EI*Length + Medium:EI + Medium:Length + (1|Replicate))“ and “lmer(Area ∼ EI*Length + Medium:EI + Medium:Length + (1|Replicate))“ for elongation index, area and length, respectively. Models were chosen based on AIC and statistical assumptions including homoscedasticity and normal distribution of residuals. Fixed variable interactions were dropped when non-significant. Predictive data was generated using the “predictInterval” function with 1000 simulations to generate overlaying box plots with 95% confidence intervals found in Fig. 1F. Libraries used: car, data.table, emmeans, lme4, merTools, readxl, performance.

**Supplementary Figure 3:**
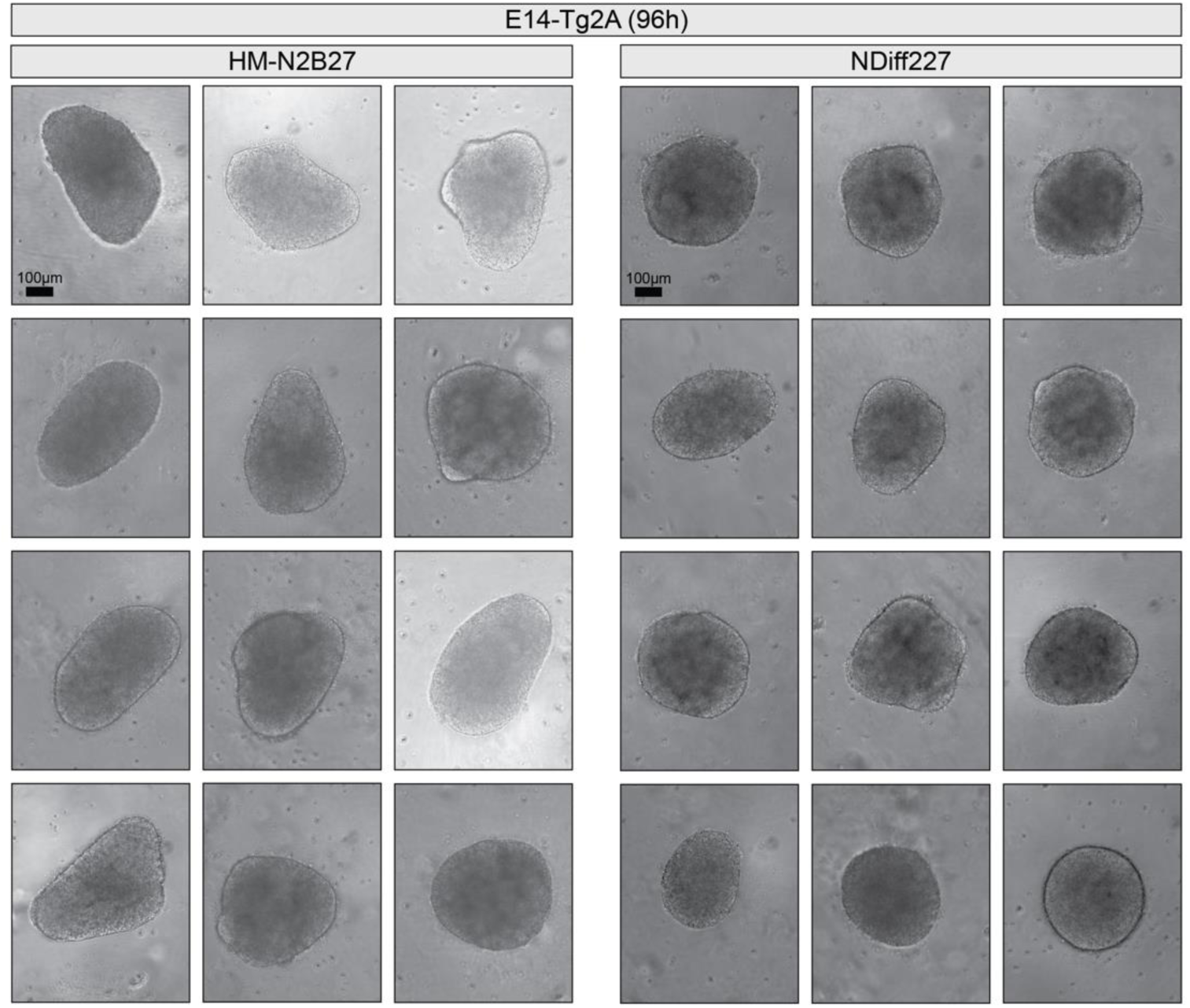
HM-N2B27 and NDiff227 gastruloids exhibit a distinct morphology at 96h. Example images from several independent experiments of E14Tg2A gastruloids cultured with either NDiff227 or HM-N2B27 media. In contrast to NDiff227 gastruloids that mostly display a round shape, the majority of HM-N2B27 gastruloids already started the elongation process at 96h. Scale bar indicates 100 μm.

**Supplementary Figure 4:**
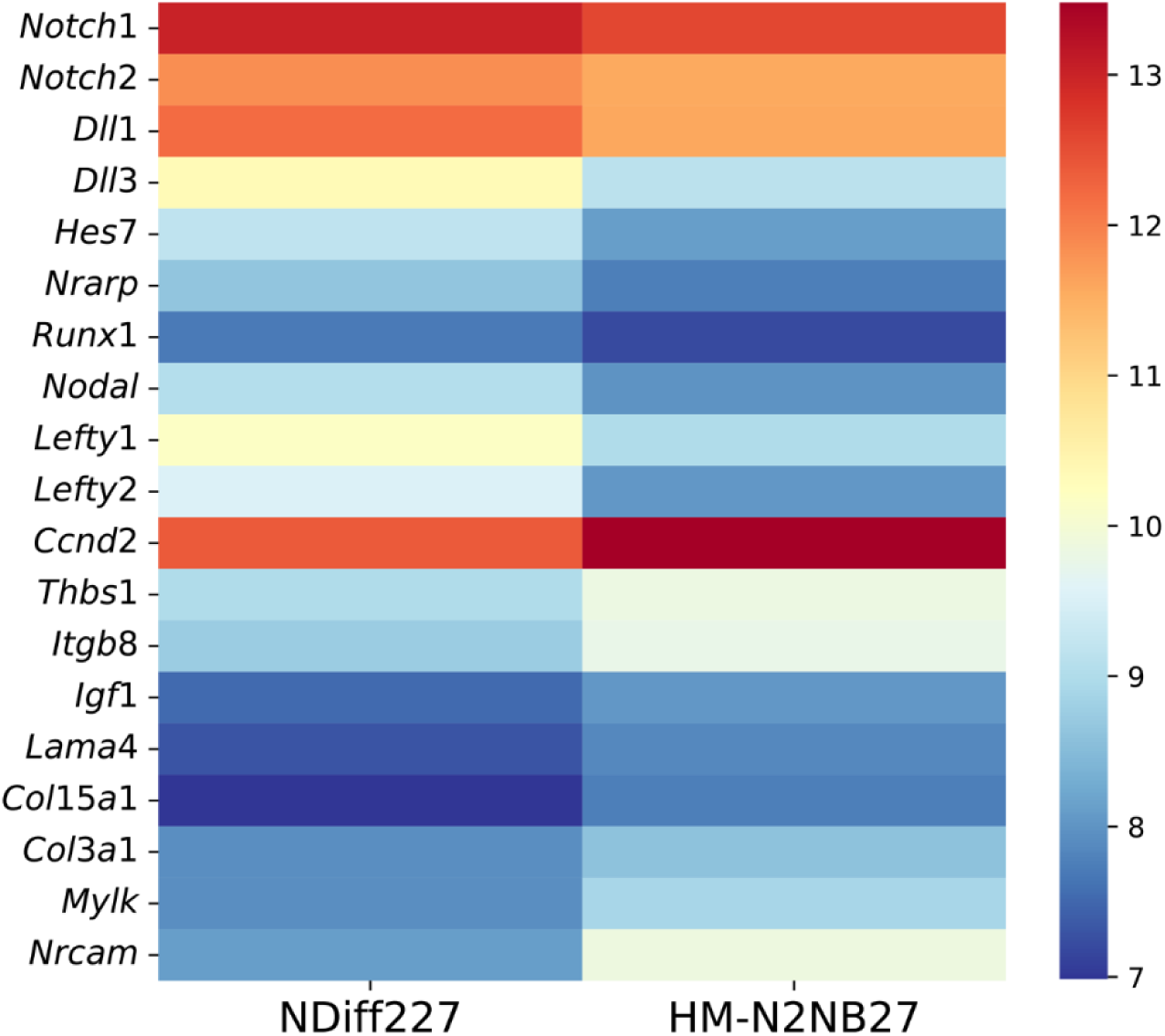
Specific transcriptomic differences between HM-N2B27 and NDiff227 gastruloids at 120h. Heatmap (mean of the replicates variance-stabilized read counts) highlighting selected genes related to the Nodal, Notch and PI3K/Akt signalling pathways that we found differently expressed between the two media conditions. Also, it is showed the expression of some genes related to biological processes that are differently enriched between HM-N2B27 and NDiff227 gastruloids (e.g. *Col3a1*, *Mylk*, *Nrcam*; Fig. 4 and **Table S5**).

## Supplemental Tables

**Supplemental Table S1:**
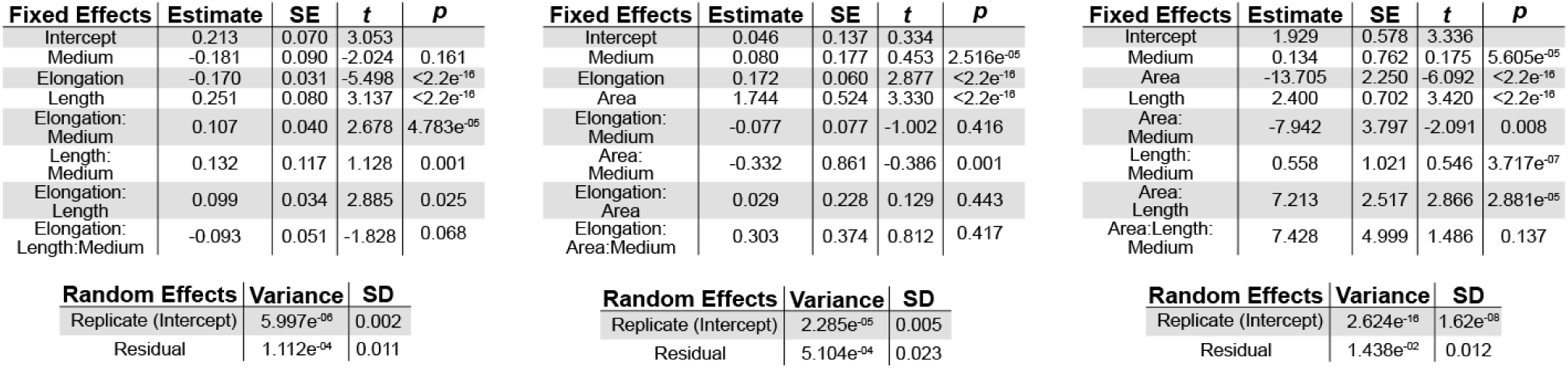
Statistical output of maximum mixed effects model. Tables include individual fixed effects variables, interactions and random effects for area (left), length (middle), and Elongation Index (right). Variables that were *p*<0.05 were dropped in refined models used for generating predictive data in **Fig 1D**.

**Supplemental Table S2:** Number of gastruloids pooled per time-point. Numbers of gastruloids in each experimental replicate for the indicated time-points, corresponding to Fig. 2. Average cell numbers from three experimental replicates ± standard deviation is shown rounded to the nearest whole number. A Two Way ANOVA followed by Bonferroni’s Post Hoc test was performed with the *p* value of the pairwise comparisons between N2B27 and NDiff227 at each time-point given to 4 decimal places. Only selected comparisons are shown for clarity.

| Time Point | No. of Gastruloids | Average Number of cells/gastruloid | p value |
| --- | --- | --- | --- |
| 48h | 96 | NDiff227: 1960 ± 65<br>HM-N2B27: 2501 ± 182 | $p = 0.1811$ (N.S) |
| 72h | 48 | NDiff227: 3397 ± 304<br>HM-N2B27: 5240 ± 312 | $p < 0.001$ (***) |
| 96h | 24 | NDiff227: 6268 ± 255<br>HM-N2B27: 11256 ± 60 | $p < 0.001$ (***) |
| 120h | 12 | NDiff227: 10810 ± 273<br>HM-N2B27: 21839 ± 311 | $p < 0.001$ (***) |

**Supplemental Table S3:** Differentially Expressed Genes shown in the volcano plot (Fig. 3A).

**Supplemental Table S4:** Top 100 genes based on loadings of PCA dimension 1 and 2 (Fig. 3B).

**Supplemental Table S5**: GO term analysis (Fig. 4).

